# Borf: Improved ORF prediction in *de-novo* assembled transcriptome annotation

**DOI:** 10.1101/2021.04.12.439551

**Authors:** Beth Signal, Tim Kahlke

## Abstract

ORF prediction in *de-novo* assembled transcriptomes is a critical step for RNA-Seq analysis and transcriptome annotation. However, current approaches do not appropriately account for factors such as strand-specificity and incompletely assembled transcripts. Strand-specific RNA-Seq libraries should produce assembled transcripts in the correct orientation, and therefore ORFs should only be annotated on the sense strand. Additionally, start site selection is more complex than appreciated as sequences upstream of the first start codon need to be correctly annotated as 5’ UTR in completely assembled transcripts, or part of the main ORF in incomplete transcripts. Both of these factors influence the accurate annotation of ORFs and therefore the transcriptome as a whole. We generated four *de-novo* transcriptome assemblies of well annotated species as a gold-standard dataset to test the impact strand specificity and start site selection have on ORF prediction in real data. Our results show that prediction of ORFs on the antisense strand in data from stranded RNA libraries results in false-positive ORFs with no or very low similarity to known proteins. In addition, we found that up to 23% of assembled transcripts had no stop codon upstream and in-frame of the first start codon, instead comprising a sequence of upstream codons. We found the optimal length cutoff of these upstream sequences to accurately classify these transcripts as either complete (upstream sequence is 5’ UTR) or 5’ incomplete (transcript is incompletely assembled and upstream sequence is part of the ORF). Here, we present Borf, the better ORF finder, specifically designed to minimise false-positive ORF prediction in stranded RNA-Seq data and improve annotation of ORF start-site prediction accuracy. Borf is written in Python3 and freely available at https://github.com/betsig/borf.

## Introduction

RNA-Sequencing (RNA-Seq) is a powerful tool to study differential gene expression, aid genome annotation and understand the biology of non-model and poorly annotated organisms (Stark et al., 2019; Salzberg, 2019; Ungaro et al., 2017). In cases where no high-quality reference genome sequence is available, reads from RNA-Seq data are commonly assembled *de-novo* to produce a reference transcriptome for downstream analysis (Eldem et al., 2017; da Fonseca et al., 2016). Several *de-novo* transcriptome assemblers are available of which Trinity (Haas et al., 2013; Grabherr et al., 2011) is currently the most commonly used (Fig. S1). A crucial step for quality control as well as functional annotation of *de-novo* transcriptome assemblies is the prediction of open reading frames (ORFs) to identify transcripts that code for protein sequences (Bryant et al., 2017).

ORF prediction in assembled transcriptome data appears to be a straightforward step. However, the identification of true coding regions can be complicated, for example, in cases where both the sense (+) and antisense (−) strand of the assembled transcript contain potential coding regions. Modern RNA-Seq assemblers use information such as sequencing design and technology used to improve assembly accuracy (Geniza and Jaiswal, 2017; Hölzer and Marz, 2019). In contrast, current ORF prediction tools disregard this information despite the obvious advantages. For example, when using RNA-Seq data originating from strand-specific libraries, which preserve the original 5’ to 3’ direction of transcripts, only coding regions predicted on the sense strand should be considered true (Zhao et al., 2015).

Another challenge in ORF prediction is the identification of the correct start site of the protein coding region. Translated ORFs generally consist of a start codon (one of the codons encoding the amino acid methionine), followed by a sequence of translatable codons that terminate at the first stop codon encountered (Kearse and Wilusz, 2017; Orr et al., 2020). Although ORFs always terminate at a stop codon, not all methionine codons are used as a translation start, and can occur within the main ORF. In addition, the start codon can be preceded by nucleotide sequences in the 5’ untranslated region (5’ UTR) which —if using the codons to predict protein sequence in the same frame as the true ORF —can appear to encode amino acids. These upstream amino acid (uAA) codons are not preceded by a start codon and therefore should not be translated. Predicting the correct translational start site from *de-novo* assembled transcripts is further complicated by the fact that assembled transcripts are not always complete (Smith-Unna et al., 2016): if a transcript is lowly expressed in the sample RNA, or if sequencing depth was too low, parts of the transcript may be missing from the final assembly. Therefore, complete transcripts can have an upstream sequence which is interpreted as the transcript assembly being ‘incomplete’, or incomplete transcripts can have an internal start codon which is interpreted as the transcript assembly being ‘complete’ leading to either the addition of uAA sequence (incorrectly extending) or the removal of ORF sequence upstream of an internal methionine (incorrectly truncating), respectively.

Transdecoder (Haas et al., 2013) is the current standard ORF predictor and annotation tool, however it does not consider these challenges. By default, Transdecoder predicts ORFs on both strands, despite strand-specific data being recommended for *de-novo* assembly (Geniza and Jaiswal, 2017). In addition, Transdecoder classifies ORFs based on their completeness, and extends the predicted ORF of all transcripts with uAAs of any length.

Here, we developed the *better ORF finder* (Borf), an ORF finder written in Python3 available at https://github.com/betsig/borf. Using Trinity *de-novo* assemblies of four model organisms as a ‘gold standard’, we show the impact that strand-specificity and correct start site identification has on ORF prediction in real data. We then tested Borf, showing that it uses suitable default parameters for *de-novo* assemblies and outperforms Transdecoder.

## Materials and Methods

### Datasets

Publicly available RNA-Seq data for four reference species —*Homo sapiens* (Chhibber et al., 2017), *Danio rerio* (Kijima et al., 2018), *Arabidopsis thaliana* (Mata-Pérez et al., 2016), and *Saccharomyces cerevisiae* (Wu et al., 2018) —were downloaded from The European Nucleotide Archive (ENA). All of the studies were run on an Illumina Hiseq2000 sequencer, resulting in 100-101bp strand-specific paired-end reads. Sample accession identifiers are given in Table S1. All samples were checked for strand-specificity using how_are_we_stranded_here (Signal and Kahlke, 2021), using the full cDNA annotations from the appropriate reference annotation (See below).

### Transcriptome assembly

All fastq files were trimmed using Trimgalore! (Babraham Bioinformatics) with default parameters. Files were concatenated together into two fastq.gz files (one for each read —R1, and R2), which was then used to assemble transcripts fully *de-novo* (i.e. not genome-guided) using Trinity (Haas et al., 2013; Grabherr et al., 2011). Default parameters were used for Trinity with the exception of –SS_lib_type RF as all samples were sequenced as fr-second strand paired end.

### Assignment of assembled transcripts to known reference transcripts

BlastN (NCBI Resource Coordinators, 2018) of Trinity.fasta against the Ensembl 98 reference cDNA database (or Araport; see Supplementary Materials) was performed (Cheng et al., 2017; Yates et al., 2020), with a maximum e-value cutoff of 1e-30. Transcripts with multiple Ensembl hits were tagged as ‘multi_hit’. To further resolve these hits, BlastX against Ensembl 98 reference.pep files was performed. The best hit was defined as the hit with lowest e-value (and if tied, highest coverage of Ensembl protein sequence, then highest number of correct peptides (p_ident * length of Blast region)). In cases where still no best hit could be identified (tied values) the hit with the lowest BlastN e-value, then the highest BlastN bitscore was taken. As Blast will produce multiple lines for the same subject and query if alignments have larger gaps, these were combined for the BlastX results by taking the maximum p ident, send, bitscore, minimum sstart, evalue, and sum of length, mismatch, gapopen.

### Translation of Trinity sequences and finding longest common sequences

All open reading frames (ORFs) in both sense (+) and antisense (−) directions were translated and the longest common subsequence (LCS; continuous, no substitutions, no gaps) to the best matching Ensembl protein sequence was chosen as the Trinity assembled transcript’s ORF. The LCS location was used to determine the transcript start site (5’ UTR length), stop site, and 3’ UTR length. Trinity sequences were classified according to the coverage of the Ensembl proteins, with those with a LCS length less than 10 being classified as ‘insufficient coverage’. The remaining translated ORFs were further classified as ‘incomplete’, ‘incomplete 3prime’, ‘incomplete 5prime’, ‘complete’, and further classed as ‘partial’ if they did not match the reference sequence exactly (See Supplementary Materials).

### Transcript expression quantification

The Trinity command align_and_estimate_abundance.pl with Kallisto (Bray et al., 2016) was used to quantify transcript expression, with raw reads aligned to the assembled Trinity transcripts.

### Coding potential of Trinity assembled transcripts

RNAsamba (Camargo et al., 2020) was used to predict coding potential scores for all Trinity assembled transcripts, using the partial length weights.

### Tool parameters and resource comparison

Both tools, Borf and Transdecoder (Haas et al., 2013), were run with default parameters on the Trinity output file (Trinity.fasta), i.e., Transdecoder was run on both sense and antisense strands. For run time comparisons, both Borf and Transdecoder.LongOrfs were run using default values on random subsets of varying sizes ranging from 2000 to 1 million transcripts from the h. sapiens Trinity assembly. Commands were timed using the unix utility ‘time’ and the ‘real’ time was used. All run time comparisons were run on a 2020 M1 Macbook Pro with 16GB of RAM.

### Data and code availability

Borf is available at https://github.com/betsig/borf (Signal, 2021a). All commands, software versions, and R scripts to reproduce our analyses are available at https://github.com/betsig/borf analysis (Signal, 2021b). Trinity assemblies and subsequent assembly annotations are available at osf.io/grpye (Signal, 2021c).

## Results

### *De-novo* assembly of model organism transcriptomes

To create a test data set for our comparisons, we performed Trinity *de-novo* assemblies of four organisms with pre-existing transcriptome annotations. Each dataset comprised 48M–210M stranded 100bp paired-end reads from an Illumina HiSeq2000 sequencer (Table S1 & S2). Trinity assembly resulted in 11,660–1,018,009 assembled genes (12,710–1,154,0004 assembled isoforms; Fig. 1A; Table S3). To ensure that transcripts were assembled with high quality, we implemented an expression filter of ≥100 read counts (typical in differential expression read filtering (Law et al., 2016)), which resulted in 7,086–131,093 transcripts (Fig. 1C; Table S3). Lowly expressed transcripts have poor ORF coverage, and removal of these transcripts resulted in higher median ORF coverage (Fig. S2). When matched to the most likely source gene using Blast (see Methods), this covered 50–70% of the respective annotated protein-coding transcriptome (Fig. 1B). We note that a large number of transcripts (>1 million in h. sapiens) had either very low expression (<10 reads), no hits to a reference cDNA sequence, or had insufficient coverage (no common subsequence ≥10AA) of a reference CDS (Fig. 1A; Table S4). We then assessed the assembled transcript’s coverage of their corresponding reference ORF, either containing the full ORF, or being incomplete at either the 5’, 3’, or both ends (see Methods). We found that the majority of assembled transcripts contain a full length ORF (Fig. 1C). Incompletely assembled transcripts on average had lost 191–263AA from their matched reference transcript ORF, or covered an average of 31-53% of the ORF (Figs 1D&E).

**Figure 1.**
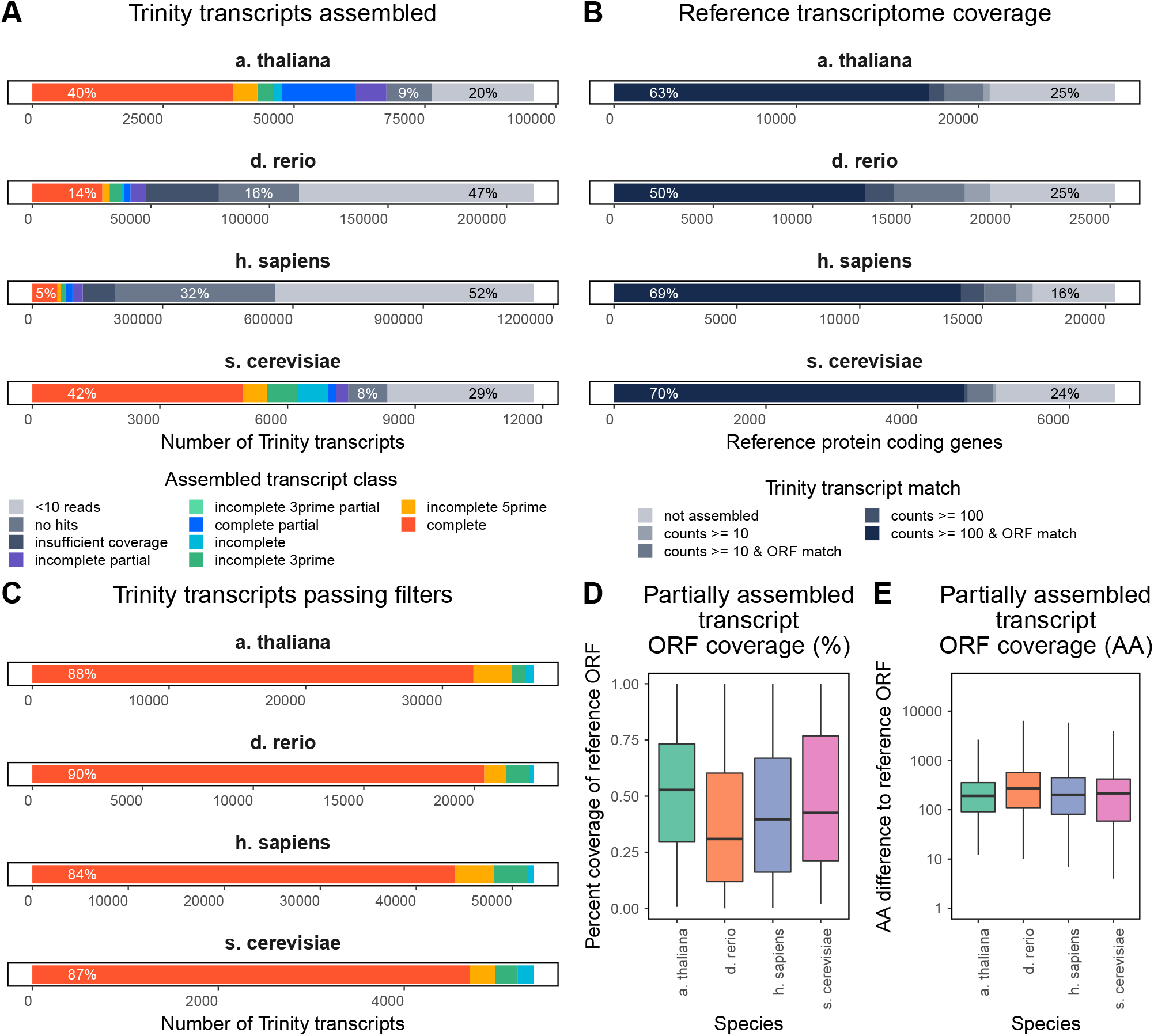
Assembly of model organism transcriptomes. A. Trinity assembled transcripts by ORF match to reference Ensembl or Araport transcripts. This covers all transcripts assembled, regardless of expression levels. B. Coverage of the Ensembl or Araport protein coding gene set. Genes are matched to either none, or at least one Trinity transcripts with at least 10 read counts. Genes were further classified by the expression and the ORF coverage of the Trinity transcript. C. Classification of Trinity assembled transcripts passing our read and longest common subsequence (LCS) filters. D. Coverage of the reference ORF in incompletely assembled transcripts E. ORF loss (AA) in incompletely assembled transcripts compared to the full reference ORF.

### Strand specificity in de-novo assemblies

Complete ORFs occur more often in transcripts with a Blast match on the sense strand (+ or sense-assembled), and antisense strand match (− or antisense-assembled) transcripts are most commonly tagged as ‘insufficient coverage’ of the ORF (LCS <10AA) suggesting that —as expected —using strand-specific data predominantly produces assembled transcripts in the correct orientation (Fig. 2A). This is supported by the expression of sense-assembled transcripts being significantly higher than antisense-assembled transcripts (Fig. 2B). The majority of these antisense-assembled transcripts were also predicted as noncoding by RNAsamba (coding probability <0.5), and may indeed be lncRNA antisense transcripts (Nevers et al., 2018; Katayama et al., 2005) (Fig. 2C). We note that there was a greater number of ‘complete’ antisense-assembled transcripts in s. cerevisiae (Fig. 2A). Given the estimated strand-specificity of the input RNA-Seq data (Table S1), we investigated the reasons for these presumably false positive ORFs being assembled and annotated at a greater relative number (Supplementary Materials). We found that this was due to + and − strand protein coding genes in the s. cerevisiae genome being in closer proximity, combined with a lack of UTR annotations which complicated correct assignment of assembled transcripts to their original gene. Extra care should therefore be taken when working with genomes of unknown gene density (as is often the case in *de-novo* RNA-Seq assemblies) to prevent accidental annotation of transcripts on the incorrect strand. We propose that predicting ORFs from strand-specific data (the typical method for library preparation in Illumina RNA-Seq (Signal and Kahlke, 2021)) should only be done on the sense strand to avoid the increase in downstream computing time, predicting ORFs that should not be translated such as antisense lncRNAs, and incorrectly annotating transcripts nearby or overlapping genes from the opposite strand.

**Figure 2.**
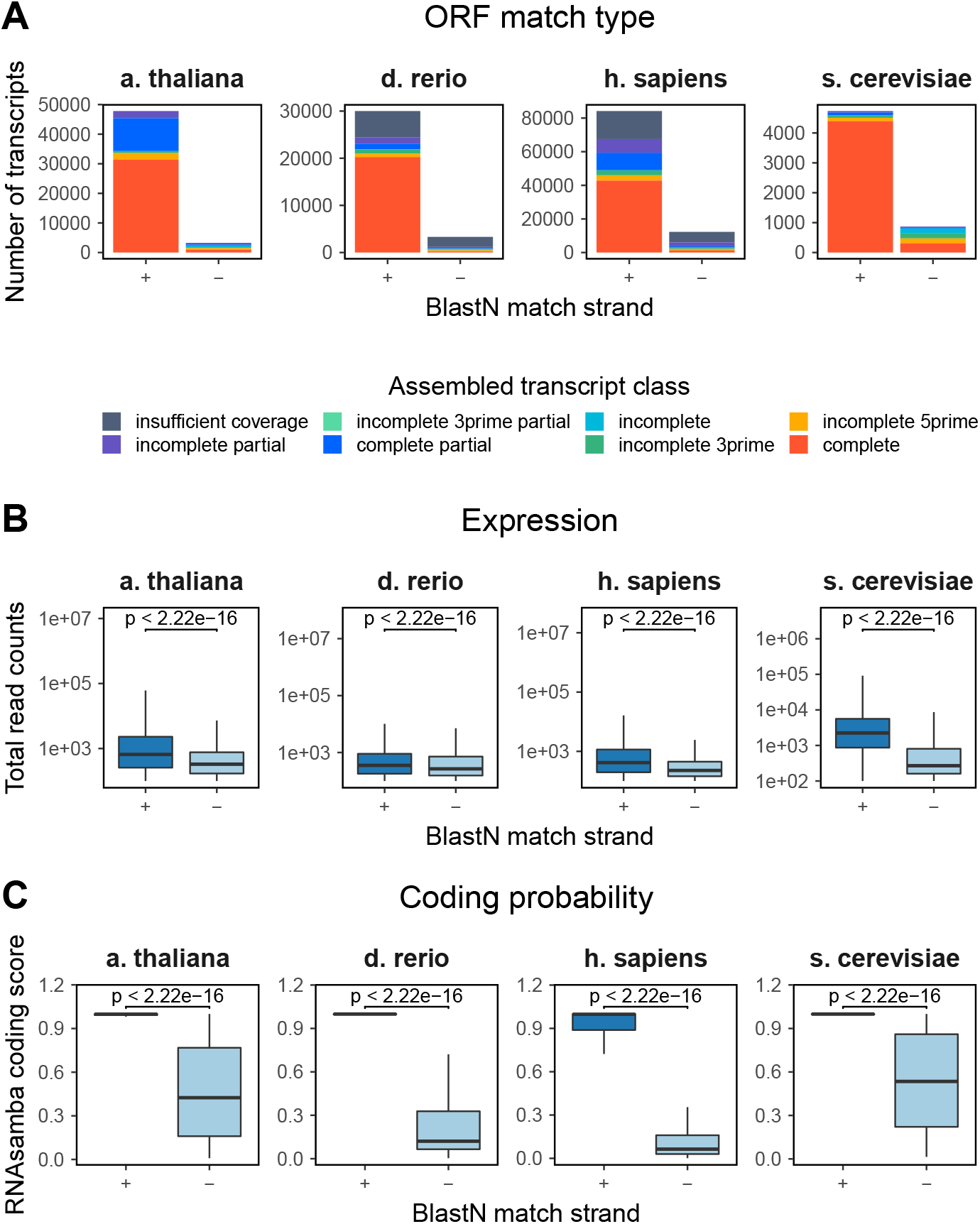
Strandedness bias in assembled transcriptomes. A. Assembled transcript ORF type of transcripts with a Blast match to the sense (+) strand or antisense (−) strand of a reference cDNA sequence. B. Read counts across all samples of transcripts with a sense or antisense strand Blast match. C. RNAsamba coding score of transcripts with a sense or antisense strand Blast match. P-values are from a Wilcoxon signed rank test.

### Uninterrupted ORF-upstream amino acids and incomplete 5’ ORFs complicate ORF prediction

Translated in-frame nucleotide sequences upstream of the ORF (ORF upstream amino acids; ORF-uAA) —although typically not translated themselves (Cross, 2016) —can complicate how the best ORF for a transcript is assigned. Presence of a stop codon upstream of a start codon indicates that the ORF can not be extended further upstream, however complications arise when no such stop codon is present. In these cases, this could mean that the transcript ORF is complete and no upstream stops occur within the 5’UTR; or the transcript ORF is incomplete and missing the 5’ end of the ORF. We found that 15–33% of transcripts had no stop codon in the ORF-uAA in reference annotations, and 11–23% of transcripts in our assembled data —which contains truly partially assembled transcripts and complete transcripts —11–23% of transcripts had no stop codon in the ORF-uAA. We then sought to find a suitable cutoff based on the assembled 5’ UTR length for accurately classifying the 5’ end of an assembled transcript as complete or incomplete. Using transcripts with complete or incomplete 5’ ends, we compared the use of ORF-uAA cutoffs from 1 to 100AA (1–300nt) in transcripts without an upstream stop on overall accuracy. For all species, in transcripts with ≥ 100 read counts, the optimal cutoff was at least 60AA (Fig. 3), and we have therefore set the default Borf cutoff at 50AA. We do however note that lower cutoffs may be more suitable to increase specificity if working with transcripts that have poor coverage and expression, as the inclusion of these increases the proportion of incompletely assembled transcripts, including those with an incomplete 5’ region, and thus affects the overall accuracy (Figs S3&S4, Table S5).

**Figure 3.**
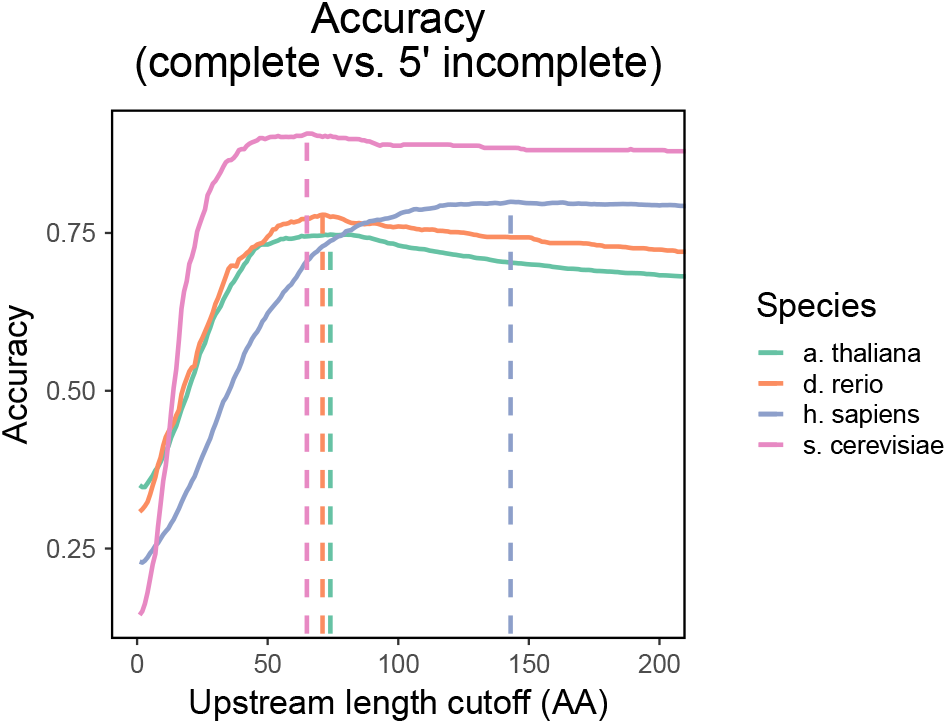
Accuracy of classifying assembled transcripts as ‘complete’ and ‘incomplete_5prime’ based on varying ORF-uAA cutoffs. For these calculations, we included transcripts with an upstream stop, and classified them as complete so we could model the relative effect the cutoff has on a whole assembly, not just the effect on transcripts without a upstream stop.

### Implementation of Borf

Borf is implemented in Python3, and can be installed using common package managers such as pip or conda. To fix the issues of strand-specificity and correct start site identification —which can appear in annotation pipelines if specific changes are not made to account for them —Borf by default returns sense strand hits only, and implements an 5’ uAA cutoff of 50 AA. Borf outputs a .pep file for amino acid sequences of each predicted ORF as well as a .txt file with data on each ORF including frame, strand, start and end sites, UTR lengths, and ORF types based on their first and last codons. In addition, we note that data strandedness can often be unknown or unclear (Signal and Kahlke, 2021), and to help prevent incorrectly using stranded or unstranded predictions on assembled datasets, Borf also implements a strandedness check on the first batch of sequences that quickly flags to users if the input data and the strand parameter do not match. Strand-specificity checks are performed by determining the strand of the longest ORF for each input nucleotide sequence, then calculating the sense strand bias (number sense strand/total ORFs). If over or under 70% of the longest ORFs are on the sense strand, Borf flags the data as likely stranded or unstranded respectively.

### Comparison of Borf to Transdecoder

Comparison of Borf and Transdecoder was performed by ORF prediction on the four reference Trinity assemblies using both tools and comparing the results to our curated annotations. Transdecoder by default performs ORF prediction on both strands of the assembled transcripts regardless of the sequenced library. Our results show that if we use the approach of taking the longest ORF for each transcript, up to 15% of transcripts incorrectly have their longest ORF on the antisense strand and either give no BlastP match, or a false positive match (Fig. 4; Table S6). If we instead use the approach of taking all ORFs, not only the longest, this not only approximately doubles the resulting AA sequences that may then need to be annotated, but also results in up to 23% false positive matches and 79% with no match to other protein sequences —due in part to a large number of antisense strand ORFs (Fig. 4; Table S7). Furthermore, Transdecoder classifies ORFs based on their completeness, and to determine if the 5’ end is incomplete, searches for any length of AA codons upstream of a start codon (M) without a stop codon. As shown above, many transcripts —both assembled and Ensembl annotated —contain uninterrupted ORF-uAAs and this approach falsely classifies and extends the predicted ORF of all such transcripts (Figs 3D & 5A). To then compare the accuracy provided by implementing a 5’ upstream AA cutoff, we took the longest Transdecoder ORF (≥100 AA) on the sense strand and compared ORF classification. Borf more often correctly assigns ORF class, except in s. cerevisiae due to negative strand bias in the assembly annotation (Table S8), increasing total accuracy by up to 12%, due to more often correctly classing ‘complete’/‘incomplete 3prime’ ORFs —even if they have a short uninterrupted ORF-uAA (Fig. 5B). Furthermore, run time of Borf is comparable to Transdecoder in small (<10000 transcripts) datasets with a median difference of +0.4 seconds, and faster in larger datasets (Fig. 5C), which are typical for *de-novo* assemblies.

**Figure 4.**
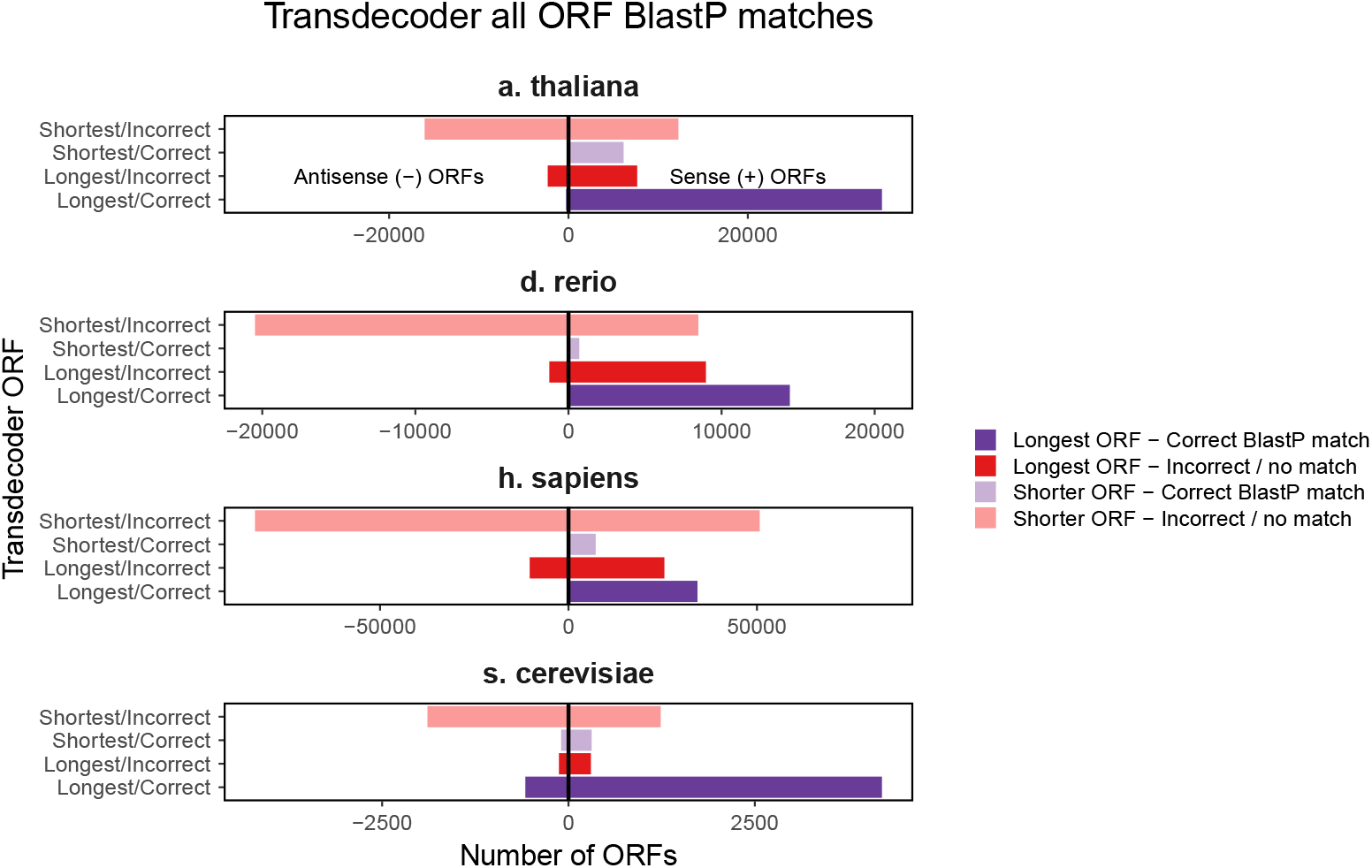
Transdecoder multiple prediction and strand accuracy. BlastP hits of the longest and shorter Transdecoder ORFs for each transcript by ORF strand.

**Figure 5.**
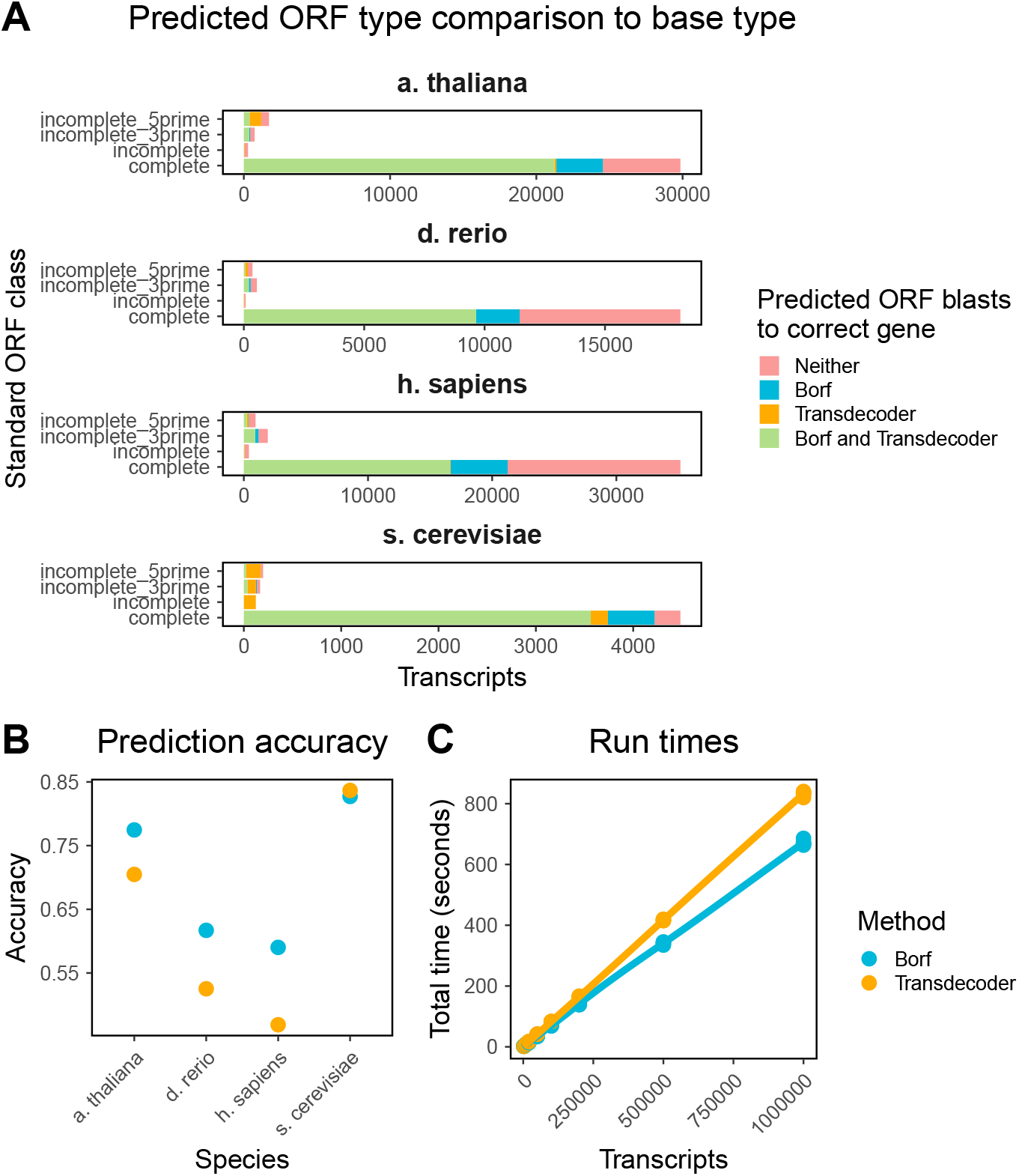
Transdecoder and Borf Performance. A. Number of transcripts in each class with correct Blast hits from each method. B. Prediction accuracy of Transdecoder and Borf. Accuracy calculated using the number of transcripts with a correctly assigned ORF classes and a correct Blast match. C. Run times for Borf and Transdecoder.LongOrfs with varying numbers of transcripts.

## Discussion

In this paper, we present Borf —an open reading frame finder for *de-novo* assembled eukaryotic transcriptome annotation. Borf was developed by taking into account several assembly characteristics to help improve ORF prediction in *de-novo* assembled transcriptome data. First, by default Borf only predicts ORFs on the sense strand. This is based on recommendations to use paired-end stranded-library sequencing for *de-novo* assemblies (Geniza and Jaiswal, 2017) as it improves the correct assembly of transcripts, particularly in more complex genomes, where it helps to correctly assemble noncoding transcripts located at the antisense strand and not confuse these with their protein coding pair. In addition, as not all *de-novo* assemblies are performed with stranded data, we added a warning if less than 70% of the longest ORFs are on the sense strand, and vice versa for running on the sense strand. This allows users to intervene and to check the strandedness of their data (through a tool such as how_are_we_stranded_here (Signal and Kahlke, 2021)) prior to any further steps. We found that including negative strand ORFs introduces false positives which do not give hits to the correct transcript or gene annotations, can leave them discarded, or annotated as ORFs of unknown function. In addition, if predicting all ORFs and not just the longest, this introduces a large number of incorrect ORF sequences, which greatly increases the time spent on annotation downstream. Second, we implemented a cutoff in classifying ORFs as 5’ incomplete or complete, and consequently reporting their ORF sequence with or without the ORF-uAA translated. Using this cutoff instead of a blanket rule where all ORFs with uAAs are classed as “5’ incomplete” allows a more accurate characterisation of protein sequences and identification of fully assembled transcripts with complete ORFs. While our default cutoff has been informed by *de-novo* assembly data and expression filtering, this can also be adjusted. We tested Borf on four Trinity *de-novo* assemblies of well annotated model organisms. While this approach was used to test across a range of genome diversity, this is not completely representative of genome and transcriptome diversity in the eukaryotic kingdom. However, patterns across all four species showed similar trends which we expect to be similar in other species, including the coverage and proportions of incomplete ORFs, and the effects of including ORF-uAAs and negative strand predictions. In conclusion, given these clear advantages, we propose Borf as the current state of the art ORF finder to be used in *de-novo* transcriptome annotation pipelines.

## Notes

### Competing Interest Statement

The authors have declared no competing interest.

