## Supplementary Materials for "Borf: Improved ORF prediction in *de-novo* assembled transcriptome annotation"

### Supplementary Methods

##### Classification of assembled ORFs

Trinity assembled sequences were classified according to the coverage of the reference proteins. For translated ORFs with a longest common subsequence (LCS) with the matched reference protein greater than 10, these were classified as (i) 'incomplete' (missing a methionine codon at the start and an asterisk at the end of the sequence), (ii) 'incomplete 3prime' (lacking only the terminating asterisk), (iii) 'incomplete 5prime' (lacking the initial methionine codon), (iv) 'complete' (includes start and stop codon and same length as the determined best hit protein in the reference database),(v) 'complete partial' (includes start and stop codon but different length as the determined best hit protein in the reference database) and (vi) 'incomplete partial' (LCS covered part of the ORF, but there was still assembled sequence upstream and downstream which did not match the ORF). In addition to the 'partial' classifications, all assembled transcripts which had assembled sequence upstream or downstream, i.e., 'incomplete 5prime' and 'incomplete 3prime' respectively, which did not match the reference ORF were tagged as 'partially assembled', and reclassified based on the extended predicted ORF (i.e. if an M was found upstream in a 'incomplete 5prime', or if a STOP was found downstream in an 'incomplete 3prime', reclassified it as 'complete') and excluded these from some analyses. For schematics of partial ORF matches and re-classifications, see Figure S5.

##### Alternative reference annotations

For s. cerevisiae, where minimal UTR annotations are included within the Ensembl annotation, the TIF-Seq dataset from (Pelechano et al., 2013) was used to annotate UTRs, requiring a count of at least 10 in at least one condition. Estimated UTR ranges were mapped to Ensembl gene locations to further improve 3' and 5' UTR ranges (i.e. excluding any annotated gene or CDS region). When genes had multiple UTRs, these were combined to give a single UTR covering the same regions. For a. thaliana, Araport 11 annotations (Cheng et al., 2017) were used instead of Ensembl annotations for all analyses. For h. sapiens, two additional altered reference transcriptomes were constructed. First, by inclusion of only transcripts with the transcript biotype "protein_coding", and second by inclusion of only 'CDS' features from the GTF annotation. These CDS regions were then used to re-annotate the parent transcript and gene boundaries in the GTF file.

### Supplementary Results

##### Factors influencing transcript assembly and annotation on the antisense strand

First, we found that the number of reads used for transcriptome assembly impacted the proportion of negative strand transcripts (Figure S6A&B), informing our use of fewer input reads for the s. cerevisiae assembly compared to h. sapiens, d. rerio, and a. thaliana to maintain comparability between each (Table S2). We also found that inclusion of noncoding regions in the reference transcriptome impacted annotation of assembled transcripts (Figure S6A). We ran our annotation protocol on the original h. sapiens assembly, but substituting the full reference transcriptome annotation for a custom annotation with only protein coding transcripts, or only the CDS regions (i.e. without UTRs) of protein coding transcripts. Removal of the UTRs for each transcript resulted in the reduction of transcripts annotated as 'complete' from 57828 to 32523, and the proportion of complete sense-assembled to complete antisense-assembled transcripts from 95% to 91%. As the s. cerevisiae reference annotation does not contain UTRs, this is likely a contributor to the observed difference in ORF match strands and types. Further, we found that genomic distance between pairs of genes on opposite strands likely impacted false positive annotation in s. cerevisiae by increasing the number of assembled transcripts which overlap in an antisense fashion - and therefore have both sense and antisense Blast matches to separate genes (Figure S7). Our annotation pipeline was conceived to primarily assign assembled transcripts to the correct isoform of a gene, and thus when using antisense hits, will select the parent transcript with the best Blast match (see Methods) and ignore strand-specificity. We did this to stop any strand biases in our annotations, and generate an annotation as close to standard non-strand-specific pipelines on stranded data. We found that upon re-annotation of the antisense-assembled transcripts with their next-best sense strand hit, s. cerevisiae ORF match types and their Blast match strands more closely resembled other species (Figure S8).

We also performed annotation using the sense strand hits only, which resulted in a 1–4.5% loss of overall protein coding genes covered by the assembly (Table S9) from a 8–25% reduction in total annotated transcripts, showing that a large number of antisense-assembled transcripts have already been assembled and annotated by a sense-assembled transcript. Together, these findings show that genome structure — in particular shorter distances between genes and smaller number of genes — can influence assembly and annotation.

### Supplementary Figures

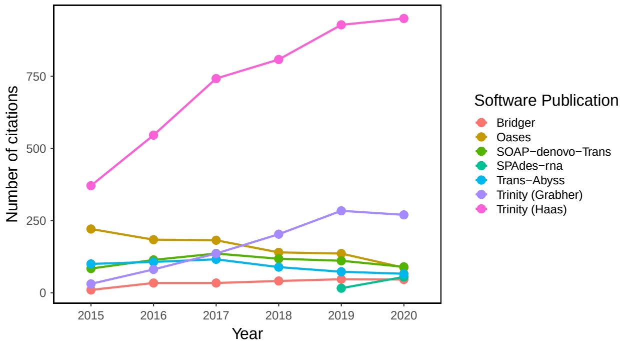


**Figure S1. Number of citations to *de novo* transcriptome assembly software.** Number of citations to the main publication (Chang et al., 2015; Schulz et al., 2012; Xie et al., 2014; Bushmanova et al., 2019; Robertson et al., 2010; Haas et al., 2013; Grabherr et al., 2011) for each transcriptome assembly software on Google Scholar. Data retrieved March 22 2021.

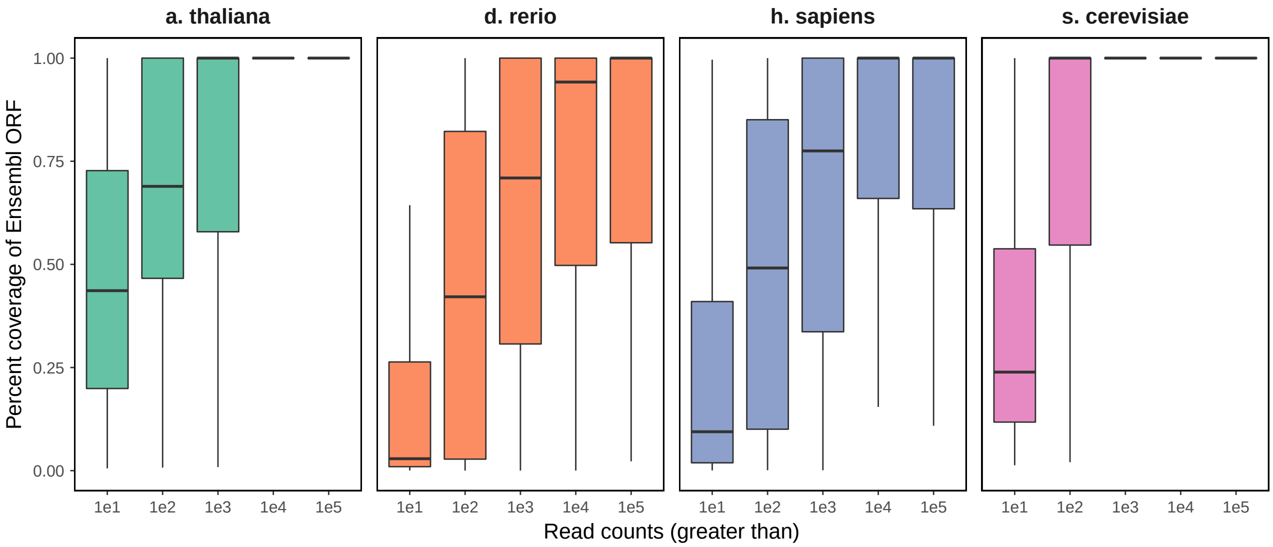


**Figure S2. *De novo* assembled transcript coverage by kallisto read counts.**

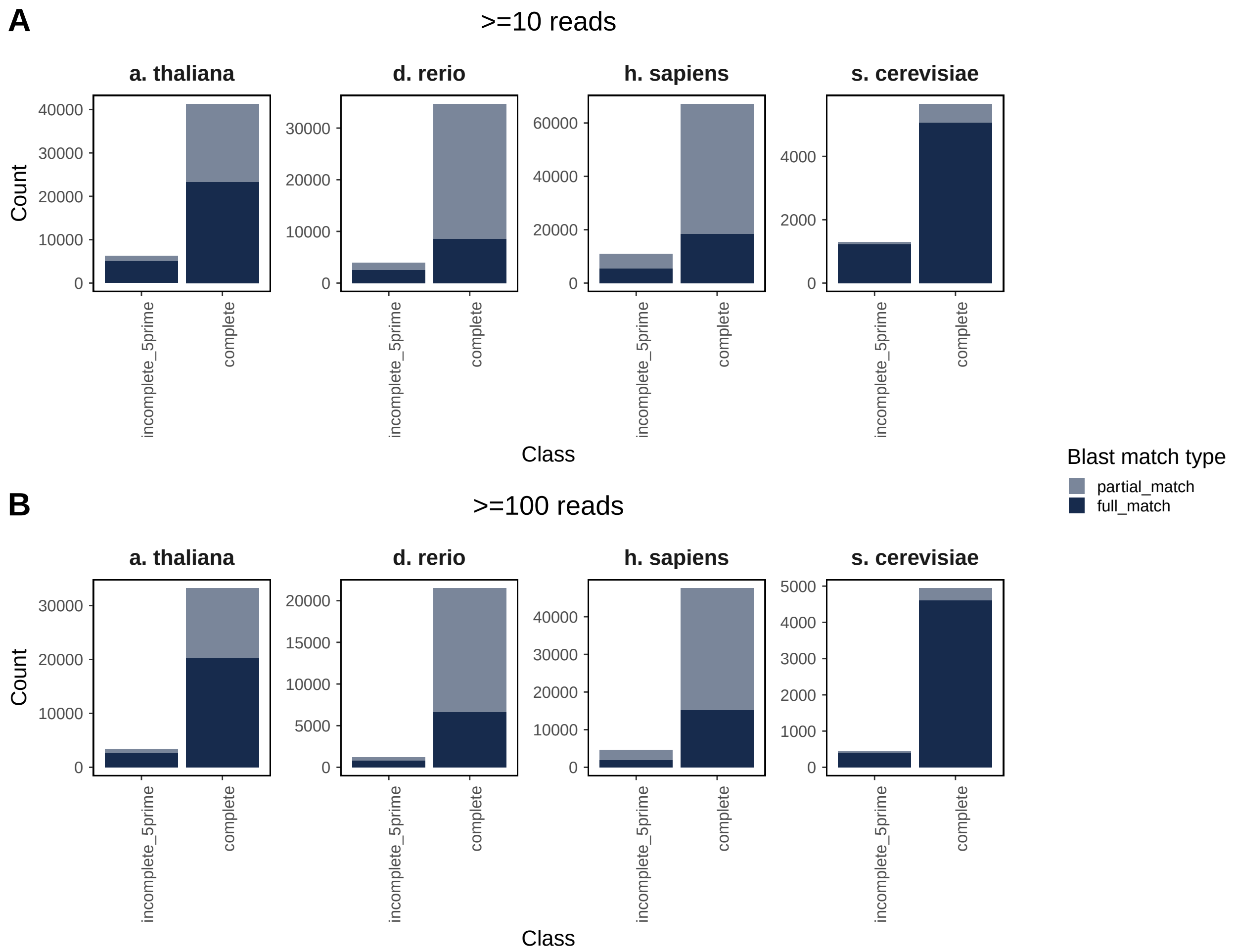


**Figure S3. Number of assembled transcripts with complete or incomplete 5' ends.** A. Transcripts with at least 10 reads. B. Transcripts with at least 100 reads. Transcripts assigned as either having complete 5' ends when 'complete' or 'incomplete_3prime', and as having incomplete 5' ends when 'incomplete' or 'incomplete_5prime'. Transcripts coloured by if the Blast match to the parent ORF was an exact full match, or a partial match.

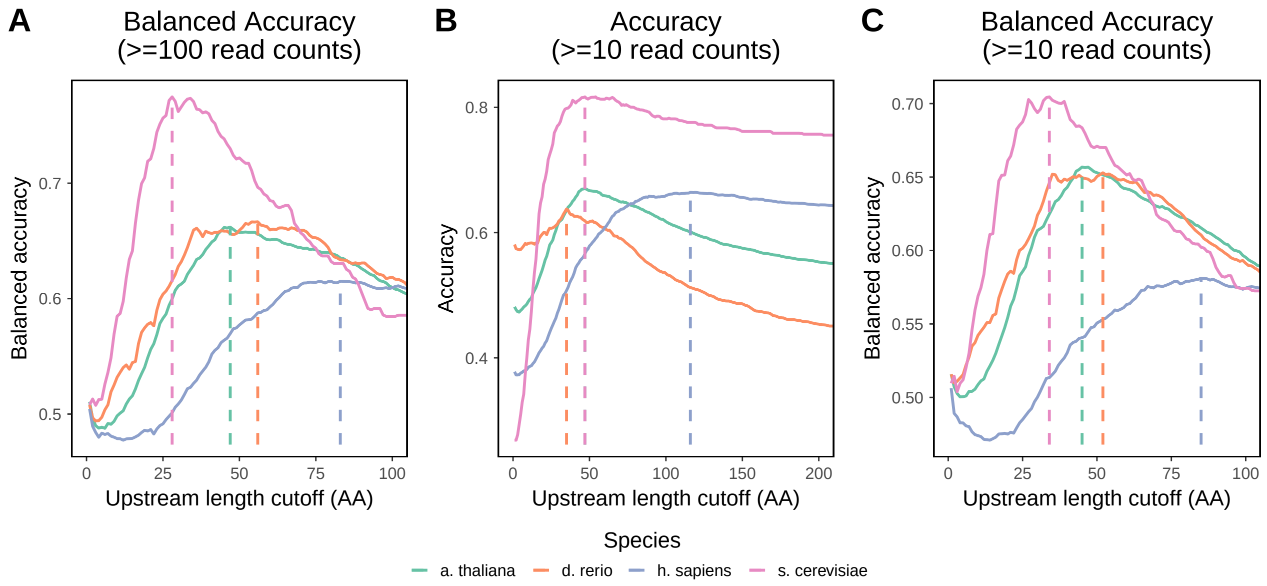


**Figure S4. Accuracy of classifying assembled transcripts as 'complete' and 'incomplete_5prime' based on varying upstream AA cutoffs.** A. Balanced accuracy in transcripts with at least 100 read counts. B. Accuracy in transcripts with at least 10 read counts. C. Balanced accuracy in transcripts with at least 10 read counts. Max accuracy or balanced accuracy is indicated by a vertical dashed line.

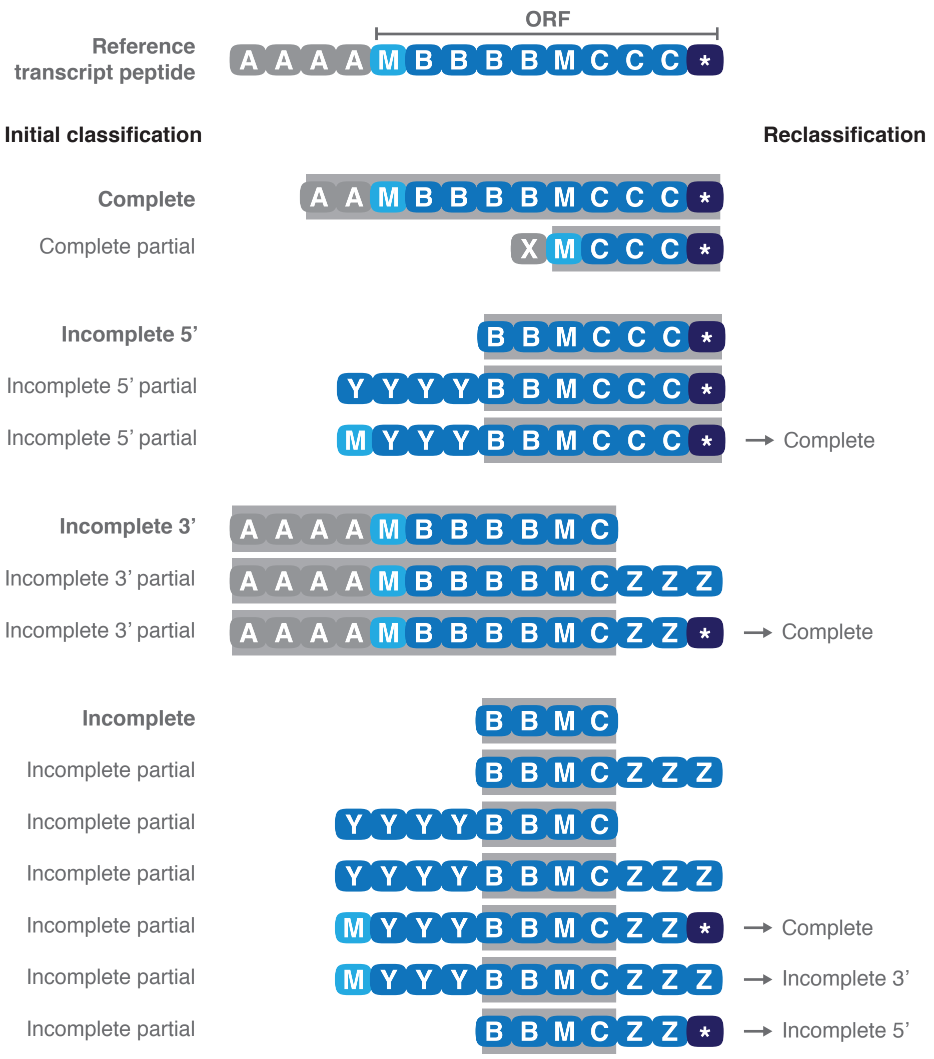


**Figure S5. Schematic of classification of assembled transcripts based on the LCS with the reference transcript.** Initial classification of assembled transcripts is given on the left, with partial matches containing some sequence which does not match the reference ORF. When alternative start or stop codons are found, these partial matches are reclassified based on presence of a start and/or stop codon (right). Blue: Open reading frame (ORF) of translated transcripts, light blue: start codon, dark blue: stop codon, grey: 5’ UTR, grey boxed area: longest common subsequence with the reference transcript.

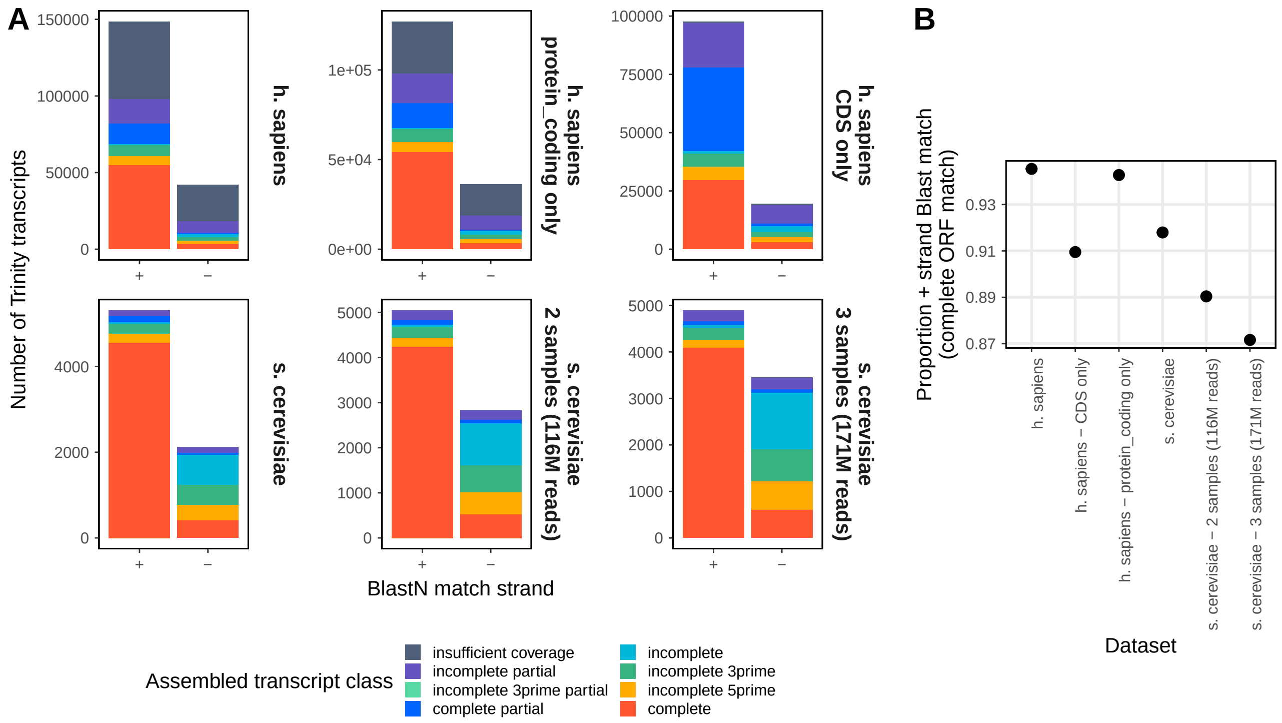


**Figure S6. Proportion of sense (+) strand annotated ORFs in h. sapiens and s. cerevisiae alternative datasets.** A. Trinity assembled transcripts by ORF match to reference Ensembl transcripts. For h. sapiens, the same Trinity assembly was re-annotated using either a reference transcriptome with only transcripts with a protein_coding transcript biotype, or only the CDS regions of these protein_coding transcripts. For s. cerevisiae, three separate assemblies were performed using one, two, or three of the RNA-Seq samples. B. Proportion of 'complete' transcripts on the sense (+) strand compared to 'complete' transcripts on the antisense (-) strand.

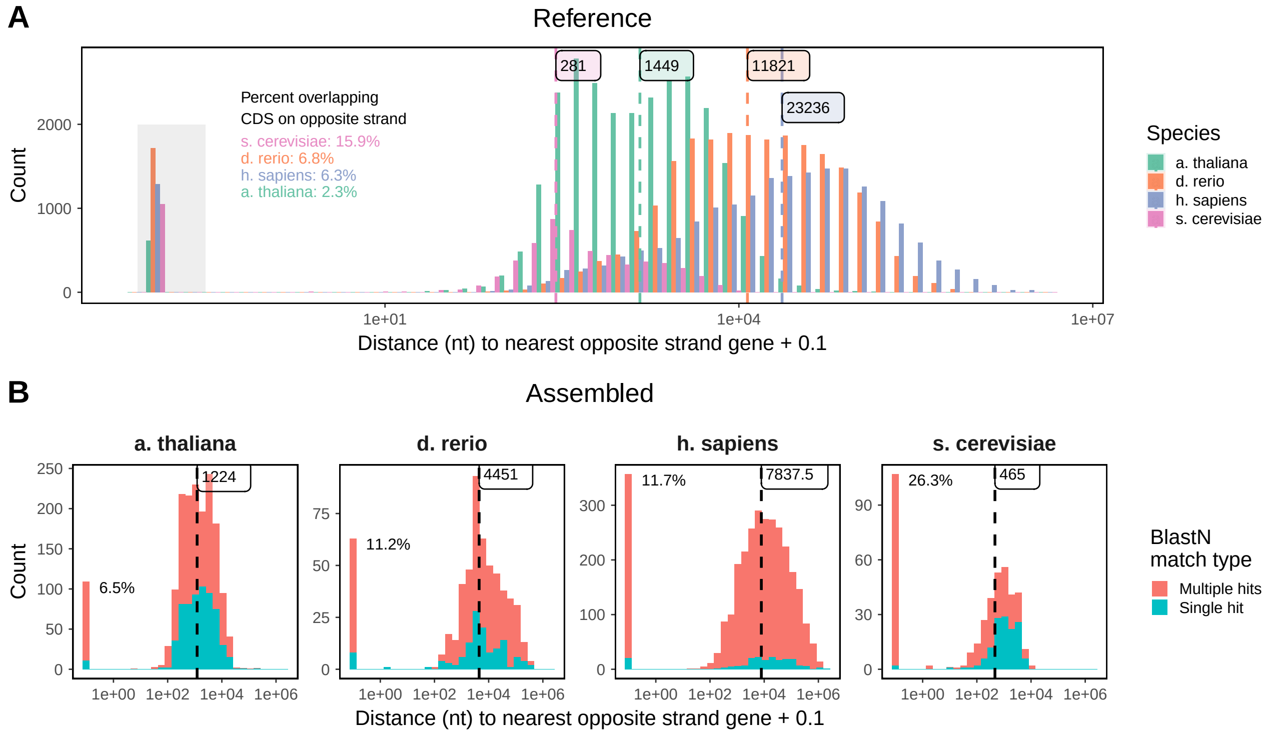


**Figure S7.** **Distances between gene CDS regions on opposite strands.** A. Distance to the nearest opposite strand gene CDS for each reference protein coding gene. Genes overlapping a CDS on the opposite strand have a distance equal to zero, are highlighted by a grey box, and have the percentage this comprises of total transcripts annotated to the right. Median distances are shown with dashed vertical lines. B. Distance from each negative strand match assembled transcript to the nearest opposite strand gene CDS in the reference transcriptome. Transcripts are shown as having either a single or multiple BlastN hits. Percentage overlapping are annotated by the first bar. Median distances are shown with dashed vertical lines.

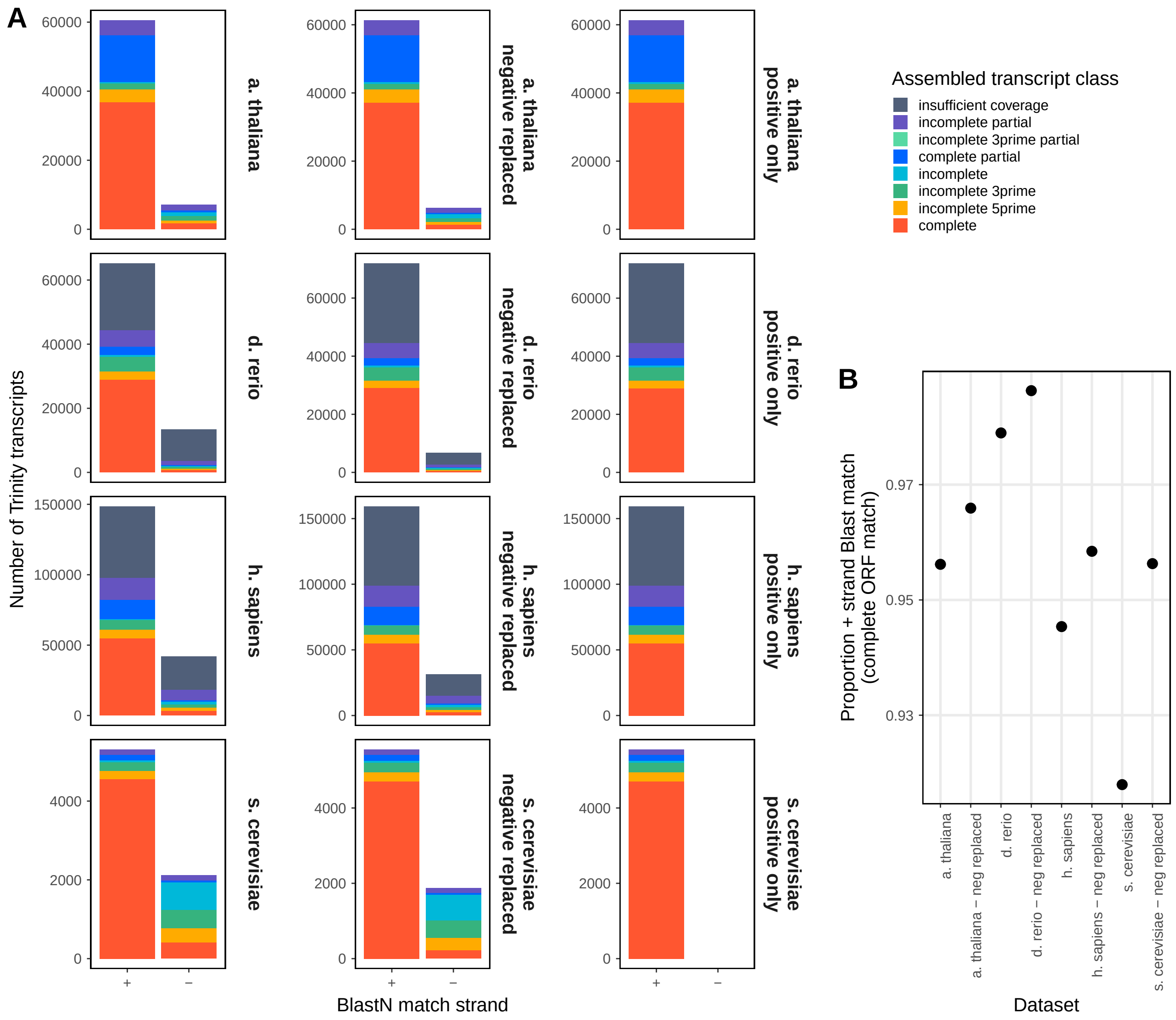


**Figure S8.** **Proportion of sense strand annotated ORFs in datasets with antisense strand hits replaced or excluded.** A. Trinity assembled transcripts by ORF match to reference Ensembl or Araport transcripts. Negative replaced datasets have all antisense BlastN match strand transcripts replaced by a sense strand annotation if possible. Positive only datasets were annotated using only sense strand BlastN matches. B. Proportion of 'complete' transcripts on the sense (+) strand compared to 'complete' transcripts on the antisense (-) strand.
